# Post-Influenza Environment Reduces *Aspergillus fumigatus* Conidia Clearance and Facilitates Invasive Aspergillosis *In Vivo*

**DOI:** 10.1101/2022.10.14.512336

**Authors:** Ko-Wei Liu, Madeleine S. Grau, Jane T. Jones, Xi Wang, Elisa M. Vesely, Matthew R. James, Cecilia Gutierrez-Perez, Robert A. Cramer, Joshua J. Obar

## Abstract

*Aspergillus fumigatus* is a human fungal pathogen that is most often avirulent in immune competent individuals because the innate immune system is efficient at eliminating fungal conidia. However, recent clinical observations have shown that severe Influenza A virus (IAV) infection can lead to secondary *A. fumigatus* infections with high mortality. Little is currently known about how IAV infection alters the innate antifungal immune response. Here, we established a murine model of IAV-induced *A. fumigatus* (IAV-Af) superinfection by inoculating mice with IAV followed 6 days later by *A. fumigatus* conidia challenge. We observed increased mortality in the IAV-Af superinfected mice compared to mice challenged with either IAV or A. fumigatus alone. *A. fumigatus* conidia were able to germinate and establish a biofilm in the lungs of the IAV-Af superinfection group, which was not seen following fungal challenge alone. While we did not observe any differences in inflammatory cell recruitment in the IAV-Af superinfection group compared to single infection controls, we observed defects in Aspergillus conidial uptake and killing by both neutrophils and monocytes after IAV infection. pHrodo-Zymosan and CM-H2DCFDA staining, indicators of phagolysosome maturation and ROS production, respectively, revealed that the fungal killing defect was due in part to reduced phagolysosome maturation. Collectively, our data demonstrate that the ability of neutrophils and monocytes to kill and clear *Aspergillus* conidia is strongly reduced in the pulmonary environment of an IAV-infected lung, which leads to Invasive Pulmonary Aspergillosis and increased overall mortality in our mouse model recapitulating what is observed clinically in humans.

**IMPORTANCE:** Influenza A virus (IAV) is a common respiratory virus that causes seasonal illness in humans, but can cause pandemics and severe infection in certain patients. Since the emergence of the 2009 H1N1 pandemic strains, there has be an increase in clinical reports of IAV infected patients in the intensive care unit (ICU) developing secondary pulmonary aspergillosis. These cases of flu-*Aspergillus* superinfections are associated with worse clinical outcomes than secondary bacterial infections in the setting of IAV. To date, we have a limited understanding of the cause(s) of secondary fungal infections in immune competent hosts. IAV-induced modulation of cytokine production and innate immune cellular function generates a unique immune environment in the lung, which could make the host vulnerable to a secondary fungal infection. Our work shows that defects in phagolysosome maturation in neutrophils and monocytes after IAV infection impairs the ability of these cells to kill *A. fumigatus* thus leading to increased fungal germination and growth and subsequent invasive aspergillosis. Our work lays a foundation for future mechanistic studies examining the exact immune modulatory events occurring in the respiratory tract after viral infection leading to secondary fungal infections.

## INTRODUCTION

*Aspergillus fumigatus* is a filamentous fungus that can be commonly found in the environment. In the immune competent host, hundreds to thousands of *Aspergillus* conidia can be inhaled every day without disease development. The innate immune system, including macrophages, neutrophils, monocytes, and dendritic cells (DCs) recognizes fungal conidia through pattern recognition receptors (PRRs) leading to their elimination through phagocytosis and killing via reactive oxygen species (ROS) and ROS independent mechanisms (1). However, when the immune system is impaired, inhaled conidia germinate, become pathogenic, and contribute to multiple diseases collectively termed aspergillosis. Chronic pulmonary colonization with *A. fumigatus* can lead to allergic bronchopulmonary aspergillosis (ABPA) most commonly found in cystic fibrosis (CF) or chronic obstructive pulmonary disorder (COPD) individuals (2, 3). Conversely, patients with severe impairments in innate immunity (ex: Chronic Granulomatous Disease (CGD), prolonged steroid treatment, or neutropenia), an acute exposure to *A. fumigatus* can lead to fungal germination and growth into hyphae resulting in biofilm formation, penetration of the lung parenchyma, and systemic dissemination (4–6).

Recent clinical reports indicate that patients admitted to the intensive care unit (ICU) due to severe Influenza virus infections may develop secondary fungal infections (7–10). Severe Influenza virus infection is now considered a major risk factor for the development of Influenza-associated invasive aspergillosis (IAPA) (11). However, the mechanistic causes leading to IAPA, especially in immune competent patients, remain poorly understood. The connections between a post-Influenza lung environment and susceptibility to secondary bacterial infection have been well documented (12, 13). In that setting, type I and type II interferons (IFN) induced by IAV infection are known to enhance susceptibility to secondary bacterial infection (14). Furthermore, the post-Influenza environment also affects innate immune cell accumulation and function within the lungs. Specifically, neutrophils and macrophages in the post-Influenza environment have decreased phagocytosis and clearance of bacterial pathogens (15, 16). Taken together these results demonstrate that innate immune responses observed in the post-Influenza environment lead to defective antibacterial innate immune clearance which can drive vulnerability to secondary bacterial infections, but the involvement of alterations in the susceptivity to secondary fungal infection remains unresolved. Recent results from a murine model of IAPA suggest that the elevated IFN production post-IAV infection induces STAT1 signaling and inhibits neutrophil recruitment, which leads IAPA (17). However, a comprehensive study examining innate immune cell function and antifungal immunity following IAV infection is needed to define the cause(s) of IAPA development in these IAV infected hosts.

In our current study, we established a murine IAV-*A. fumigatus* (IAV-*Af*) superinfection model that we used to examine the modulation of the antifungal innate immune response within the post-IAV lung environment. As expected, mice from the IAV-*Af* superinfection group exhibited higher morbidity and mortality, fungal biofilm formation, and increased fungal burden in the lungs. We further observe that normal inflammatory immune cell recruitment occurs in the post-IAV lung environment, yet there is a significant impairment in fungal phagocytosis and killing *in vivo*. The defective killing of *A. fumigatus* observed in pulmonary neutrophils and monocytes after IAV infection is due in part to defective phagolysosome maturation. Therefore, our findings demonstrate that modulation of neutrophil and monocyte function in a post-IAV environment contributes to defective fungal clearance, leading to disease development and eventual mortality of the host.

## RESULTS

### Influenza A virus (IAV) infection increases susceptibility to invasive pulmonary aspergillosis in mice

To understand why Influenza infected immune competent hosts are more susceptible to IPA, we developed an *IAV-Af* superinfection model with immune competent mice. C57BL/6J mice were first challenged intranasally with either PBS or a sub-lethal dose of IAV PR/8/34 H1N1 (100 EID50). Six days later, mice were challenged oropharyngeally with either PBS or 3.4 × 10^7^ *A. fumigatus* (CEA10) conidia for an additional 8-36 hours (Fig. 1A). C57BL/6J mice first inoculated with IAV and then inoculated with *A. fumigatus* (IAV-*Af* superinfection) had significantly more mortality compared to mice challenged with IAV only or *A. fumigatus* only. In the IAV-*Af* superinfection group, mice succumbed to infection as early as 1-day post *A. fumigatus* inoculation, with 100% mortality observed by 5 days post-fungal challenge (Fig. 1B).

**Fig. 1.**
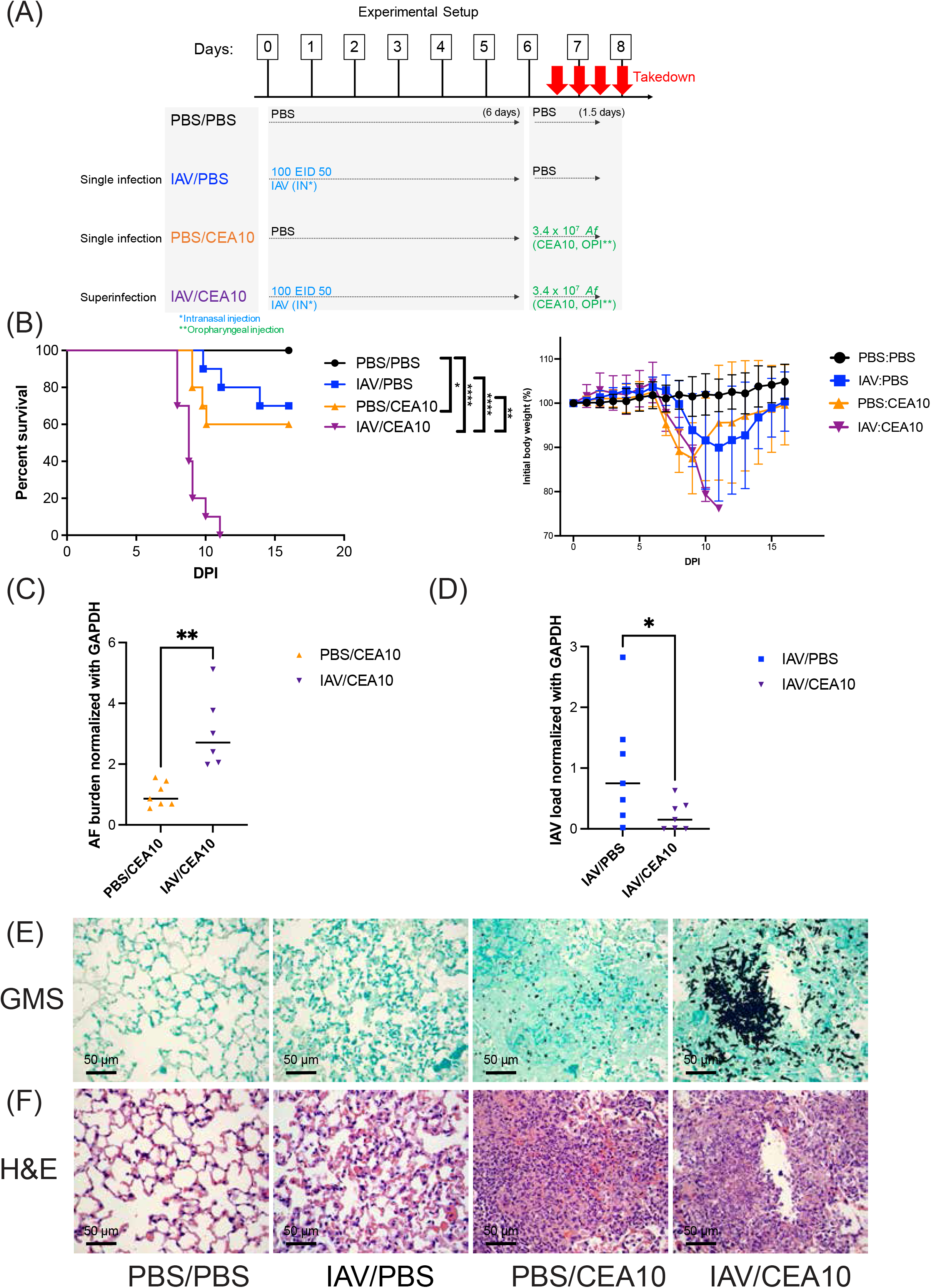
Influenza A virus infection aggravates IA disease progression. (A) Schematic design of IAV-AF infection mouse model. C57BL/6 mice were inoculated with 100 EID 50 IAV or PBS at Day 0 followed by 3.4*10^7^ CEA10 conidia or PBS at Day 6. Mice were euthanized at either 8, 24, 36, 48 hours post CEA10 inoculation. (B) Survival curve (left) of mice with PBS inoculation, IAV single infection, CEA10 single infection and IAV/CEA10 superinfection (n=10) and the weights (right) of mice in each group. Two independent experiments were performed, and data are shown as the representative results. (C) For quantification of pathogen load, RNA was isolated from the lungs of mice exposed to either 6 days of IAV or PBS followed by 36 hours of CEA10 or PBS inoculation. Fungal burden was examined by quantitative RT-PCR on *A. fumigatus* 18s rRNA (n=7). (D) Viral load was examined by quantitative RT-PCR on viral matrix protein (n=7). (C)-(D) are representative of three independent experiments. (E) For lung histology, mice were euthanized after 6 days of IAV or PBS incubation followed by 48 hours post CEA10 or PBS inoculation. Representative histology images of mice lungs were observed with GMS staining and (F) H&E staining. Two independent experiments were performed with n=5 per experiment. The Log-rank test and Gehan-Breslow-Wilcoxon test were performed for statistical analysis of the survival curve and nonparametric analyses were performed (Mann-Whitney, single comparisons) for the pathogen load. All error bars represent standard deviations. NS P>0.05; * P≤0.05; ** P≤0.01; *** P≤0.001; **** P≤0.0001.

Increased mortality coincided with higher fungal burden in the lungs of the IAV-*Af* superinfected group compared to the *A. fumigatus* only infection group (Fig. 1C). Based on our experimental design, most of the IAV would be cleared by the host at the time we collected the lung samples for viral load (Day 7.5 post IAV infection), but, interestingly, the quantification of the Influenza viral load by qRT-PCR showed even lower virus load in the IAV-*Af* superinfection group compared to IAV single infection group, indicating functional antiviral immunity in the superinfection group (Fig. 1D). To assess the development of IAPA in our mouse model, we examined lung sections with GMS and H&E staining for fungal burden and immune cell recruitment, respectively. While most of the inoculated *A. fumigatus* remained as conidia at 2-days post-inoculation in the *A. fumigatus* only infection group, in the IAV-*Af* superinfection group we observed that *A. fumigatus* conidia had germinated significantly, even forming biofilms at the infection foci (Fig. 1E). These data demonstrate that the respiratory environment found after IAV infection results in defective restriction of *A. fumigatus* germination which correlated with the development of IAPA in our IAV-*Af* superinfection model.

### Post-Influenza immunity does not affect immune cell recruitment during infection with the highly virulent *A. fumigatus* CEA10 strain

Host innate immunity, which is largely mediated by neutrophils, macrophages and dendritic cells play important roles in preventing IPA (18–23). Quantitative deficiency in any of these immune cells can limit fungal recognition and/or inhibition of conidial germination and fungal clearance. H&E staining indicates that substantial inflammatory immune cell accumulation occurs in the infection site of both the *A. fumigatus* only and IAV-*Af* superinfection groups, suggesting a robust cellular innate immune response is occurring even in the post-IAV lung environment (Fig. 1F). Since, a previous study indicated STAT1-dependent inhibition of neutrophil recruitment in the post Influenza environment (17), we next asked whether there were any differences in composition and/or absolute number of the infiltrating immune cells in our IAV-*Af* superinfection model. We leveraged multiple immune staining panels (Table S1, Fig. S1-S3) for flow cytometry analysis to examine the lung cellularity in our IAV-*Af* superinfection model (24). At 36 hours post-*A. fumigatus* challenge, we observed equivalent numbers of total lung cells, neutrophil (Ly6G^(+)^ CD11b^(+)^), monocyte (CD11b^(high)^ CD64^(-)^ MHCII^(-)^) and type 2 conventional DC (cDC2) (CD103^(-)^ CD11b^(+)^) in both the *A. fumigatus* only infection and IAV-*Af* superinfection groups that was increased compared to the PBS control and IAV single infection groups (Fig. 2A, 2B, 2E and 2G). We also observed equivalent increases in interstitial macrophages (CD11b^(high)^ CD64^(+)^) and plasmacytoid DC (pDC) (CD317^(+)^) in both the IAV only infection and IAV-*Af* superinfection groups compared to PBS control (Fig. 2D and 2H). We also examined the cellular composition by calculating the percentage of each cell type (Fig. S4). Similar to cell number observations, both the *A. fumigatus* only infection and IAV-*Af* superinfection groups had a significant increase in the percentage of neutrophils in the leukocyte population compared to PBS controls and there was no difference between those two experimental groups (Fig. S4A). We observe an increased percentage of interstitial macrophages and pDC populations in the IAV only infection group compared to the *A. fumigatus* only infection and IAV-*Af* superinfection groups though there was no difference in total cell number (Fig. S4C and S4G). Collectively, our flow cytometry analysis suggests that the recruitment of multiple immune cell types to the lung following IAV infection and *A. fumigatus* infection still contribute to corresponsive cell recruitment in the IAV-*Af* superinfection environment.

**Fig 2.**
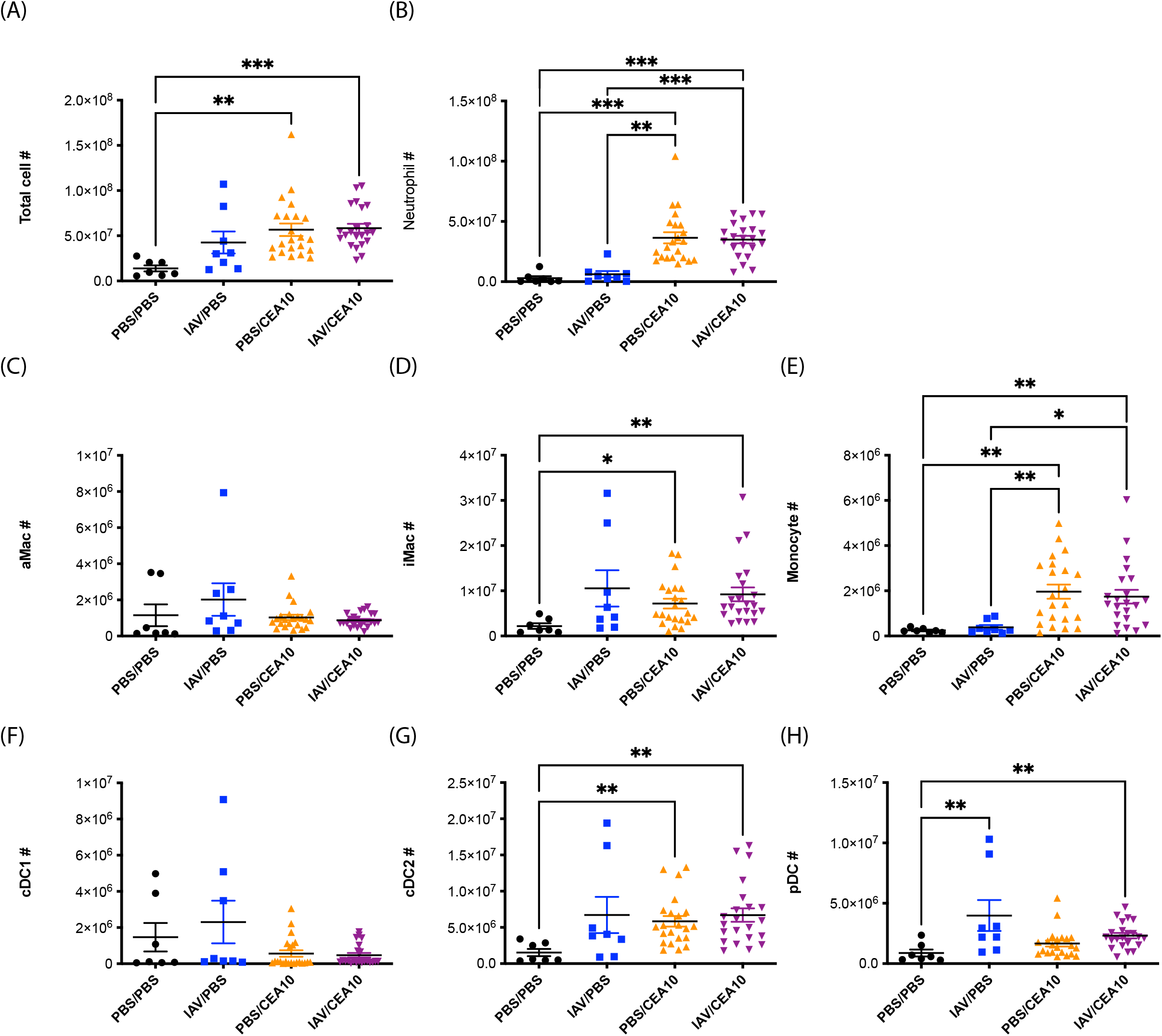
Influenza A virus infection does not affect lung cellularity during IA. C57BL/6 mice were inoculated with 100 EID 50 IAV or PBS at Day 0 followed by 3.4*10^7^ CEA10 conidia or PBS at Day 6. Mice were euthanized at 36 hours post CEA10 or PBS inoculation for lung cellularity experiments. Three independent experiments were performed, and data are shown as the combined results (PBS/PBS group: n=7; IAV/PBS group: n=8; PBS/CEA10 group: n=22; IAV/CEA10 group: n=22). All lung cell numbers were acquired by flow cytometry as indicated: (A) total lung cells, (B) neutrophils (CD45^(+)^Ly6G^(+)^CD11b^(+)^), (C) alveolar macrophages (Ly6G^(-)^CD103^(-)^SiglecF^(+)^CD11b^(+)^), (D) interstitial macrophages (Ly6G^(-)^CD103^(-)^SiglecF^(-)^CD11b^(hi)^CD64^(+)^), (E) monocytes (Ly6G^(-)^CD103^(-)^SiglecF^(-)^CD11b^(hi)^CD64^(-)^MHCII^(-)^), (F) CD103^(+)^cDC1 (MHCII^(+)^CD11c^(+)^CD11b^(-)^CD103^(+)^), (G) CD11b^(+)^ cDC2 (MHCII^(hi)^CD11c^(hi)^CD103^(-)^CD11b^(+)^), (H) pDC (MHCII^(+)^CD11c^(+)^CD11b^(-)^CD103^(-)^CD317^(+)^). Kruskal-Wallis with Dunn’s multiple comparisons were performed for statistical analyses. All error bars represent standard deviations. NS P>0.05; * P≤0.05; ** P≤0.01; *** P≤0.001; **** P≤0.0001.

### Neutrophils and monocytes in the post-Influenza lung environment have defects in antifungal killing mechanisms

Since we did not observe any differences in either the absolute number of innate immune cells or their composition, we hypothesized that the lung environment found after IAV infection results in altered antifungal effector function(s) within the recruited innate immune cells. This seemed likely, since defects in innate immune cell function, rather than recruitment, can lead to IPA development in corticosteroid treated patients and mice (25, 26). In order to test this hypothesis, we utilized the fluorescent *Aspergillus* reporter (FLARE) assay to quantify both the uptake and viability of *A. fumigatus* conidia within specific immune cell types in the murine lungs (Fig. S5) (33). Thirty-six hours post-inoculation of AF633-labeled mRFP-CEA10 (FLARE) conidia, both neutrophils (Fig. 3A) and monocytes (Fig. 3B) showed a slight decrease in conidial uptake with an almost two-fold increase in conidial viability within the IAV-*Af* superinfection group compared to *A. fumigatus* only infection. In contrast, we observed no differences in conidial uptake or viability within alveolar macrophages (Fig. 3C) or interstitial macrophages (Fig. 3D). Within cDC2 cells, we observed a very minor decrease in conidial uptake, but no change in conidial viability (Fig. 3E). Besides the increase in intracellular viability of *A. fumigatus* in neutrophils and monocytes, we also noticed that the remaining extracellular, free conidia showed higher viability in the single cell suspension from murine lungs in the FLARE assay (Fig. 3G). In parallel, single cell lung suspensions showed an almost two-fold increase in fungal CFUs from the IAV-*Af* superinfection group when compared to the *A. fumigatus* only infection group (Fig. 3F), which corroborates our observations with the FLARE assay. Collectively, these data show that both neutrophils and monocytes have defects in their antifungal phagocytotic and killing processes in the post-IAV environment, which could explain why *A. fumigatus* can germinate, grow, and form biofilms in the post-IAV lung environment ultimately leading to the development of disease.

**Fig. 3.**
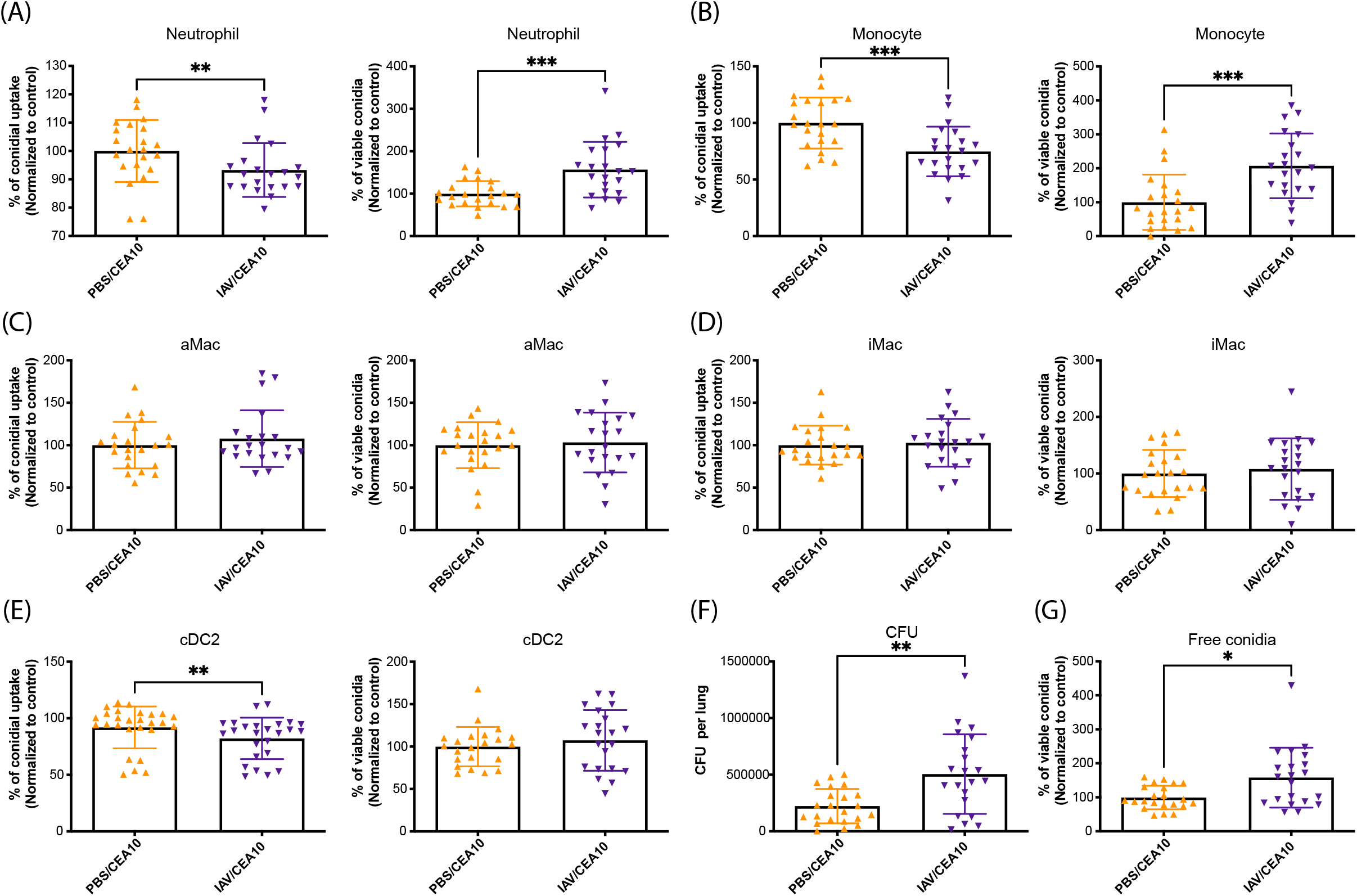
Defects in leukocyte mediated fungal killing post Influenza A virus infection. C57BL/6 mice were inoculated with 100 EID 50 IAV or PBS at Day 0 followed by 3.4*10^7^ FLARE (mRFP^(+)^/AF633^(+)^) conidia or PBS at Day 6. Mice were euthanized at 36 hours post FLARE conidia or PBS inoculation. The percentage of cells positive for conidial tracer (AF633^(+)^) and conidial viability within the immune cells (mRFP^(+)^/AF633^(+)^) were analyzed. Phagocytosis and conidial viability were examined in (A) neutrophils (CD45^(+)^Ly6G^(+)^CD11b^(+)^), (B) monocytes (Ly6G^(-)^CD103^(-)^SiglecF^(-)^CD11b^(hi)^CD64^(-)^MHCII^(-)^), (C) alveolar macrophages (Ly6G^(-)^CD103^(-)^SiglecF^(+)^CD11b^(+)^), (D) interstitial macrophages (Ly6G^(-)^CD103^(-)^SiglecF^(-)^CD11b^(hi)^CD64^(+)^), and (E) CD11b^(+)^ cDC2 (MHCII^(hi)^CD11c^(hi)^CD103^(-)^CD11b^(+)^). (F) The viability of FLARE conidia within immune cells in the lung suspension was assessed by colony forming units (CFUs). (G) The viability of free FLARE conidia in the lung suspension was shown as the percentage of mRFP^(+)^cells in free conidia population (Alexa633^(+)^FSC^(low)^SSC^(low)^). Three independent experiments were performed, and data are shown as the combined results (PBS/CEA10 group: n=22; IAV/CEA10 group: n=21). Mann-Whitney, with single comparisons were performed. All error bars represent standard deviations. NS P>0.05; * P≤0.05; ** P≤0.01; *** P≤0.001; **** P≤0.0001.

### Neutrophils and monocytes induce ROS production normally in the post-IAV lung environment

ROS production by innate leukocytes play an important role in controlling pathogen infections, particularly *Staphylococcus aureus* and *A. fumigatus*, as highlighted by patients with X-linked chronic granulomatous disease (27, 28) and mice lacking NADPH oxidase components (29). Previous studies have demonstrated that impaired ROS production by innate immune cells is associated with decreased fungal killing, even when their phagocytotic function (30). Therefore, we hypothesized that neutrophils and monocytes found in the post-IAV lung environment were impaired in their ability to induce ROS production. To test this hypothesis in using the IAV-*Af* superinfection model, we isolated immune cells from infected mice and incubated them with the cellular dye CM-H2DCFDA to detect total ROS production by both neutrophils and monocytes. We then determined both the percentage of cells with a positive signal from CM-H2DCFDA, as well as the amount of ROS produced on a per cell basis. We chose to examine the ROS response during early fungal infection since increased fungal germination and growth were observed by 24 hours post *A. fumigatus* infection. To do this we collected lung cells at 8 hours post *A. fumigatus* infection. We observed increased ROS production in neutrophils and monocytes from both *A. fumigatus* only infection and IAV-*Af* superinfection groups compared to both the PBS control and IAV only infection groups. Furthermore, we observed no decrease in ROS production from the neutrophils and monocytes from IAV-*Af* superinfection group compared to the *A. fumigatus* only infection suggesting that the fungal-induced ROS burst was not impaired in the post-IAV lung environment (Fig. 4A and 4B). For the percentage of ROS produced cells, we observed an increased percentage in IAV induced neutrophils and *A. fumigatus* induced monocytes, but there was no significant difference between IAV or *A. fumigatus* only infection and IAV-*Af* superinfection groups (Fig. S7A and S7B). Our data demonstrate that neutrophils and monocytes are still capable of significant ROS production during early *A. fumigatus* infection in the post-IAV environment indicating a defect in ROS production from these two critical innate immune cells is not likely responsible for the development of disease in our model.

**Fig. 4.**
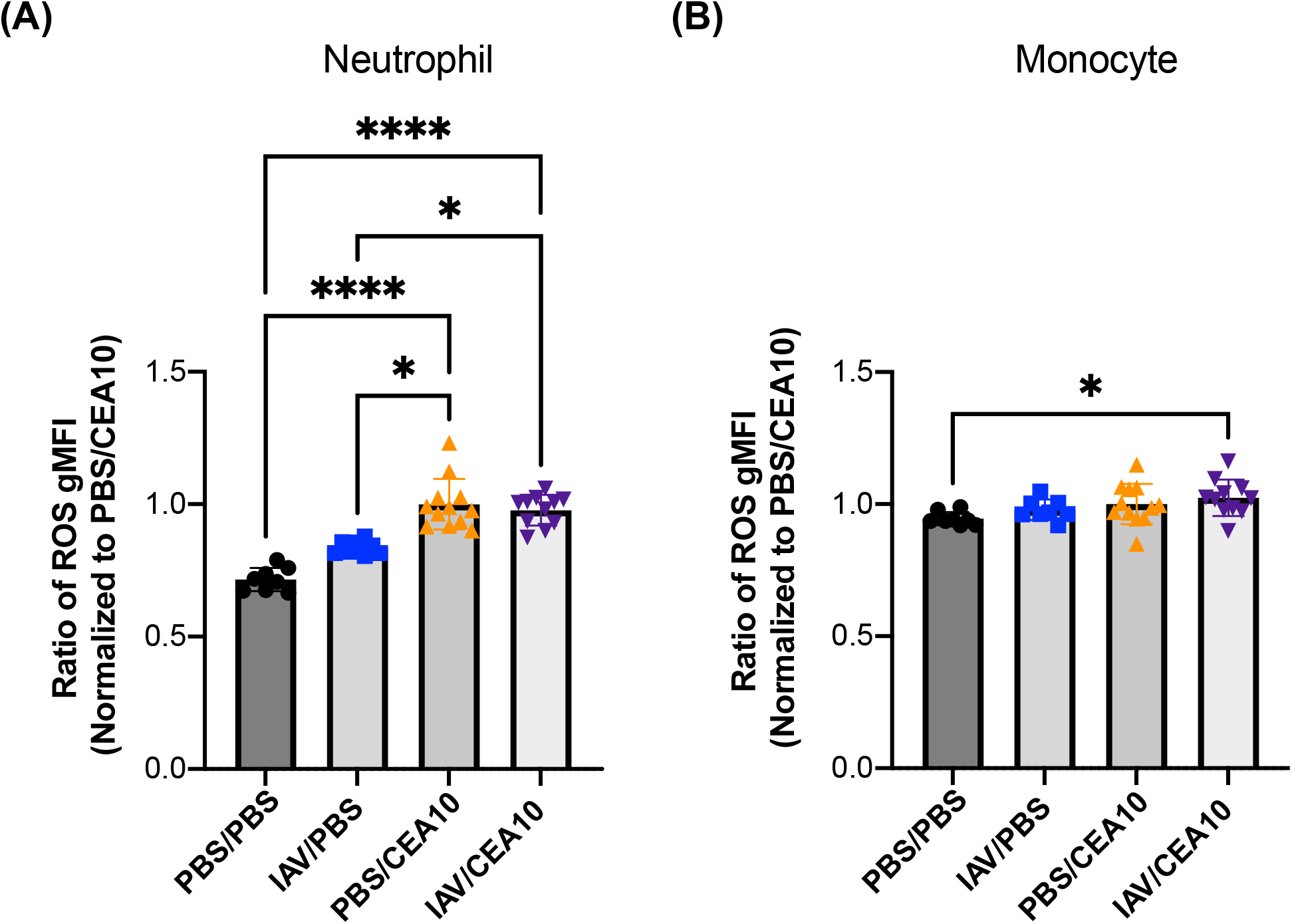
Post influenza immunity does not hinder Neutrophil or Monocyte ROS production. C57BL/6 mice were inoculated with 100 EID 50 IAV or PBS at Day 0 followed by 3.4*10^7^ CEA10 conidia or PBS at Day 6. Mice were euthanized at 8 hours post CEA10 or PBS inoculation for ROS measurement. Lung cell suspensions were stained with CM-H2DCFDA for 30 minutes and then stained for neutrophils and monocytes. ROS production was measured by the signal from CM-H2DCFDA staining in (A) neutrophils (Ly6G^(+)^) and (B) monocytes (Ly6G^(-)^SiglecF^(-)^CD11b^(hi)^CD64^(-)^MHCII^(-)^). Two independent experiments were performed, and data are shown as the combined results (PBS/PBS group: n=8; IAV/PBS group: n=8; PBS/CEA10 group: n=12; IAV/CEA10 group: n=11). Kruskal-Wallis with Dunn’s multiple comparisons was performed for statistical analyses. All error bars represent standard deviations. NS P>0.05; * P≤0.05; ** P≤0.01; *** P≤0.001; **** P≤0.0001.

### Phagolysosome maturation is impaired in neutrophils and monocytes found in the post-IAV lung environment

Rapid maturation of the phagosome through LC3-associated phagocytosis and subsequent phagosome maturation to the phagolysosome is necessary for antifungal killing and control of *A. fumigatus* germination and growth (31, 32). Since ROS production was not impaired by neutrophils and monocytes isolated from the post-IAV lung environment, we next hypothesized that defects in phagolysosome maturation within leukocytes from the IAV-*Af* superinfection contributes to the impaired antifungal killing. To test this hypothesis, we quantified phagolysosome maturation using pHrodo-Zymosan staining in the neutrophils and monocytes 8 h after *A. fumigatus* challenge. We determined the percentage of currently active cells by the percentage of cells with a positive signal from a color change of pHrodo-Zymosan and the amount of mature phagolysosomes in these active cells by their intensity of pHrodo-Zymosan signal. In our murine model, both neutrophils and monocytes from the IAV only infection group had significant reductions in their pHrodo-Zymosan signal from mature phagolysosomes compared to the PBS control (Fig. 5A and 5B). Similar to our observation in the IAV only infection group, we also observed reduced signal from mature phagolysosomes in neutrophils and monocytes from the IAV-*Af* superinfection group compared to the PBS control (Fig. 5A and 5B). These data suggest that the post-IAV lung environment significantly impacts phagolysosome maturation in neutrophils and monocytes, which correlates with the impaired intracellular killing ability against fungal conidia we observed in the IAV-*Af* superinfection group in the FLARE experiment (Fig. 3).

**Fig. 5.**
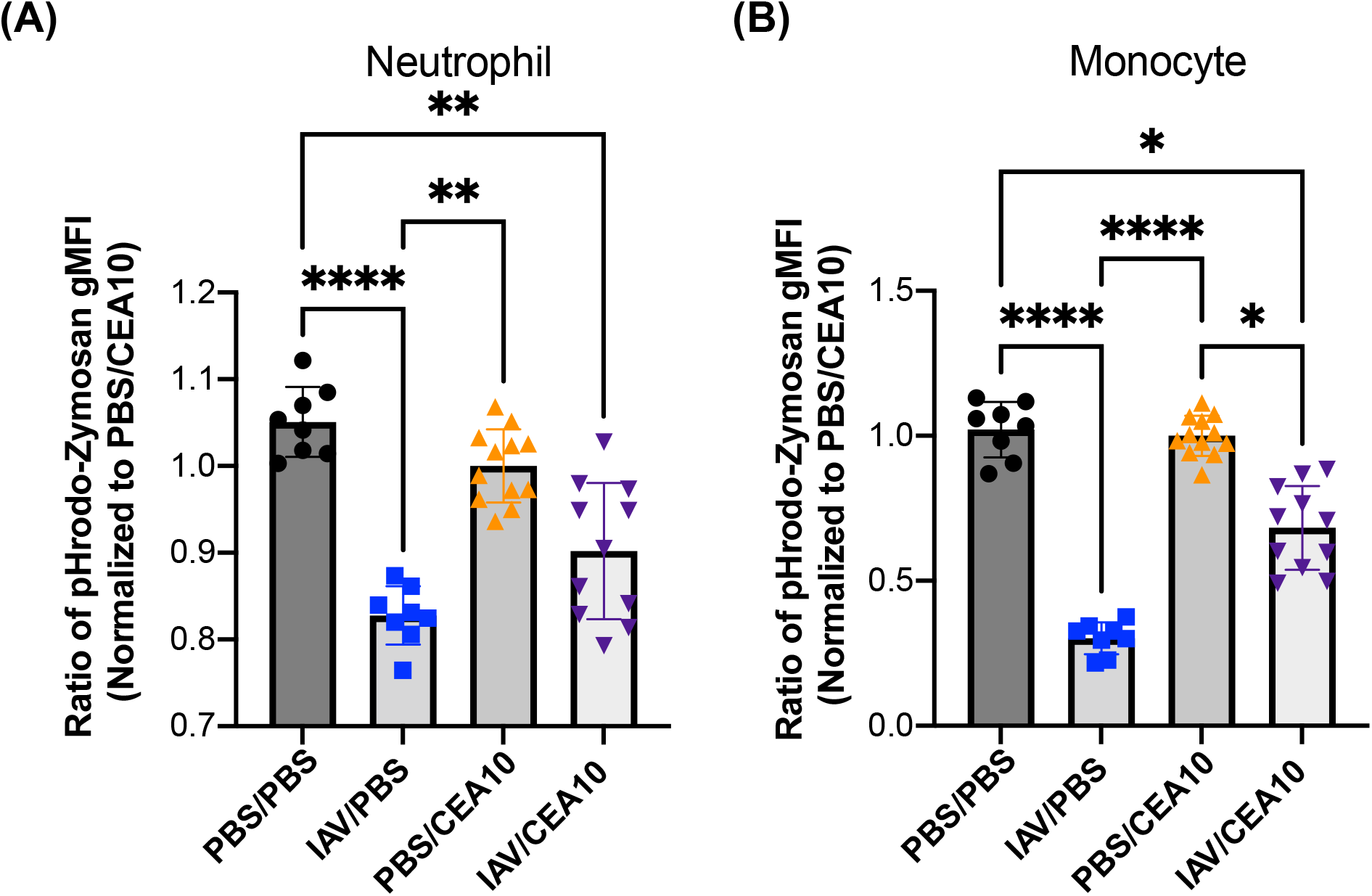
Reduced phagolysosome maturation can be detected in Neutrophils and Monocytes during early viral infection. C57BL/6 mice were inoculated with 100 EID 50 IAV or PBS at Day 0 followed by 3.4*10^7^ CEA10 conidia or PBS at Day 6. Mice were euthanized at 8 hours post CEA10 or PBS inoculation for phagolysosome maturation analysis. Lung cell suspensions were incubated with pHrodo-Zymosan for 2 hours and then stained for neutrophils and monocytes. The phagolysosome maturation level was measured by the signal from the color change of pHrodo-Zymosan in (A) neutrophils (Ly6G^(+)^) and (B) monocytes (Ly6G^(-)^SiglecF^(-)^CD11b^(hi)^CD64^(-)^MHCII^(-)^). Two independent experiments were performed, and data are shown as the combined results (PBS/PBS group: n=8; IAV/PBS group: n=8; PBS/CEA10 group: n=12; IAV/CEA10 group: n=11). Kruskal-Wallis with Dunn’s multiple comparisons was performed for statistical analyses. All error bars represent standard deviations. NS P>0.05; * P≤0.05; ** P≤0.01; *** P≤0.001; **** P≤0.0001.

To check for proportion of cells that are still capable of responding to fungal PAMPs in the post-IAV environment, we examined the percentage of cells with a mature phagolysosome. We observed that the percentage of lung neutrophils with mature phagolysosomes from *A. fumigatus* only infection and IAV-*Af* superinfection groups were reduced compared to PBS controls, but with no difference between the IAV-*Af* superinfection and either single infection groups (Fig. S8A). Intriguingly, lung monocytes from the IAV only infection and IAV-*Af* superinfection groups showed reduced mature phagolysosome containing cells in the population than both the PBS control and *A. fumigatus* only infection groups (Fig. S8B). To further test the connection between the phagolysosome maturation defect and conidial killing in the non-exhausted neutrophils from the post-IAV environment, we combined the FLARE experiment with pHrodo-Zymosan staining in our animal model. With confocal imaging, we observed Ly6G^(+)^ neutrophils containing mature phagolysosome signal as well as live or dead conidia (Fig. 6A). The pHrodo-zymosan^(+)^ neutrophils in the lung from IAV-*Af* superinfection mice suggest these neutrophils were still functional in the post-IAV environment (Fig. 6B). However, these pHrodo-zymosan^(+)^ neutrophils had less mature phagolysosome signal (Fig. 6C) and show decreased conidial killing (Fig. 6E). Collectively, these data suggest that neutrophils and monocytes in the post-IAV environment are impaired in phagolysosome maturation (Fig. 5), which correlates with decreased fungal killing (Fig. 3) and increased fungal burden (Fig. 1).

**Fig. 6.**
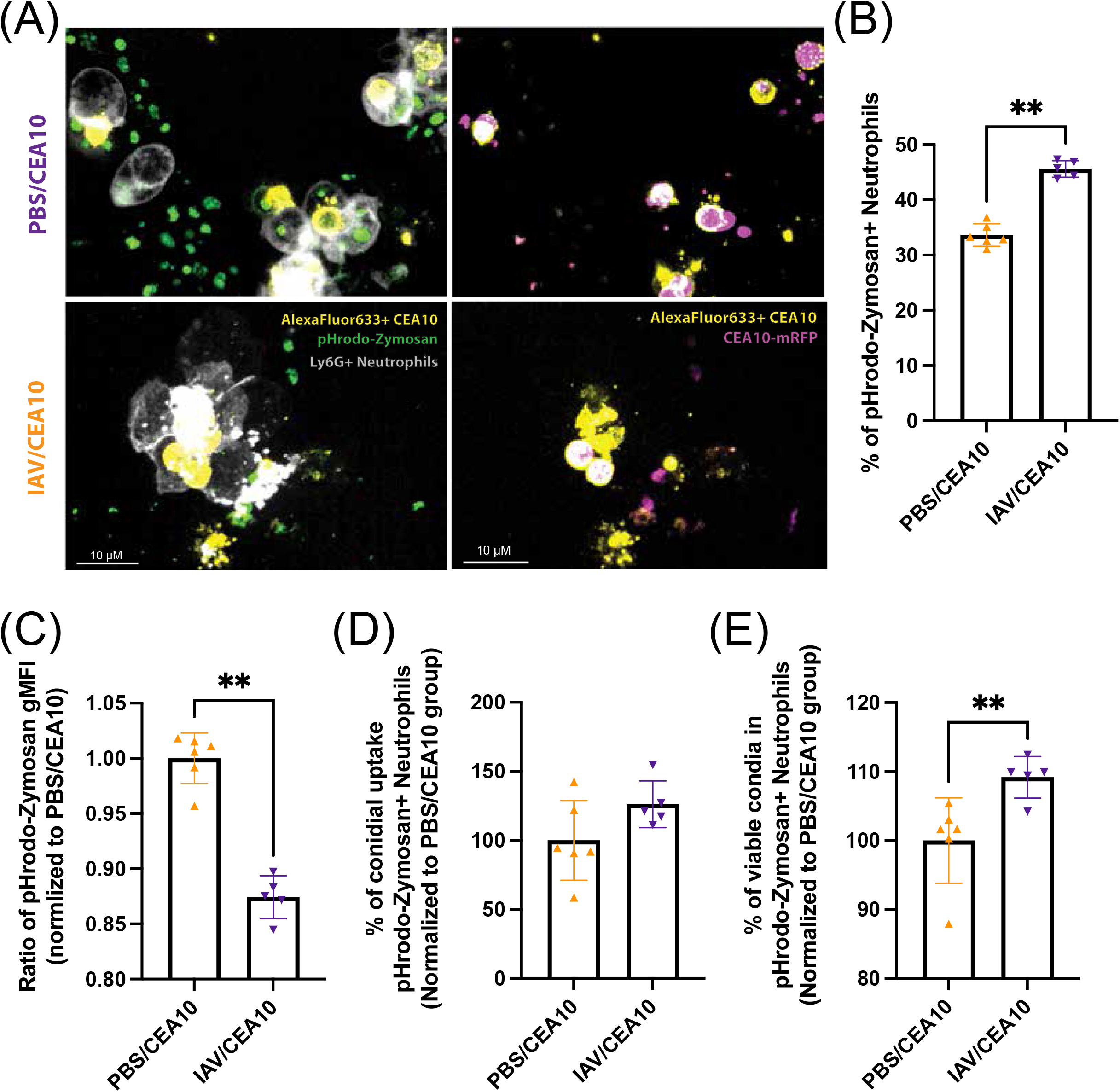
Functional neutrophils from the post-IAV environment show decreasing phagolysosome maturation and conidial killing. C57BL/6 mice were inoculated with 100 EID 50 IAV or PBS at Day 0 followed by 3.4*10^7^ FLARE (mRFP^(+)^/AF633^(+)^) conidia or PBS at Day 6. Mice were euthanized at 36 hours post FLARE conidia or PBS inoculation. Lung cell suspensions were incubated with pHrodo-Zymosan for 2 hours and then stained for neutrophils. (A) Representative confocal images of neutrophil labeling (Ly6G-Pb, white), mature phagolysosome (pHrodo-Zymosan, green), labeled conidia (Alexa633, yellow) and conidial viable marker (mRFP, pink). Right images feature labelled conidia, neutrophils, and pHrodo-Zymosan signal. Left images feature labelled conidia and RFP to indicate fungal viability. Fungal conidia with no RFP signal are considered ‘Dead’. (B) Currently functional neutrophils were indicated as pHrodo-Zymosan^(+)^ cells. (C) The amount of mature phagolysosome was shown by the gMFI of pHrodo-Zymosan in Ly6G^(+)^pHrodo-Zymosan^(+)^ cells. The percentage of cells positive for conidia (AF633^(+)^) and conidial viability within the immune cells (mRFP^(+)^/AF633^(+)^) were analyzed as (D) the percentage of conidial uptake in Ly6G^(+)^pHrodo-Zymosan^(+)^ cells and (E) the percentage of viable conidia in Ly6G^(+)^pHrodo-Zymosan^(+)^ cells. This repeated experiment was done as the combination of FLARE experiment (Figure 3) and phagolysosome maturation measurement (Figure 5) (PBS/CEA10 group: n=6; IAV/CEA10 group: n=5). Mann-Whitney, with single comparisons were performed. All error bars represent standard deviations. NS P>0.05; * P≤0.05; ** P≤0.01; *** P≤0.001; **** P≤0.0001.

### Increased inflammation but decreased PRR gene expression in the IAV-*Af* superinfection environment

In order to investigate the potential upstream pathway(s) contributing to defective phagolysosome maturation and fungal clearance we saw in the IAV-*Af* superinfection model, we performed qRT-PCR of antifungal genes in RT^2^ Profiler PCR Arrays (PAMM-147ZD) with immune cell RNA from lungs of mice challenged with *A. fumigatus* only and IAV-*Af* superinfection. At 8 hours post-*Af* challenge, we found increased mRNA levels for *Nlrp3, Pycard, Ptgs2, Cd36, Cxcl10, Nfkb1, Mapk14* and *Ccr5* and decreased mRNA levels for *Il10, Il2* and *Jun* in the IAV-*Af* superinfection group when compared to mice challenged only with *A. fumigatus* (Fig. 7). Interestingly, despite an enhanced inflammatory environment in IAV-Af superinfection conditions, which likely reflects the increased fungal growth, we observed a decrease in the mRNA levels of *Tlr9, Scarf1* and *Colec12*, as well as *Irak4* which is involved in myddosome and TLR9 signaling (Fig. 7). Our data reveal that while IAV-*Af* superinfection mice have a robust inflammatory response, there is a specific decrease in PRRs mRNA levels that could alter the early host-fungal interaction and drive the impaired phagolysosome maturation and antifungal killing observed in neutrophils and monocytes. Future studies will seek to explore the mechanism(s) underlying this viral induced defect in phagolysosome maturation in the presence of *A. fumigatus* conidia.

**Fig. 7.**
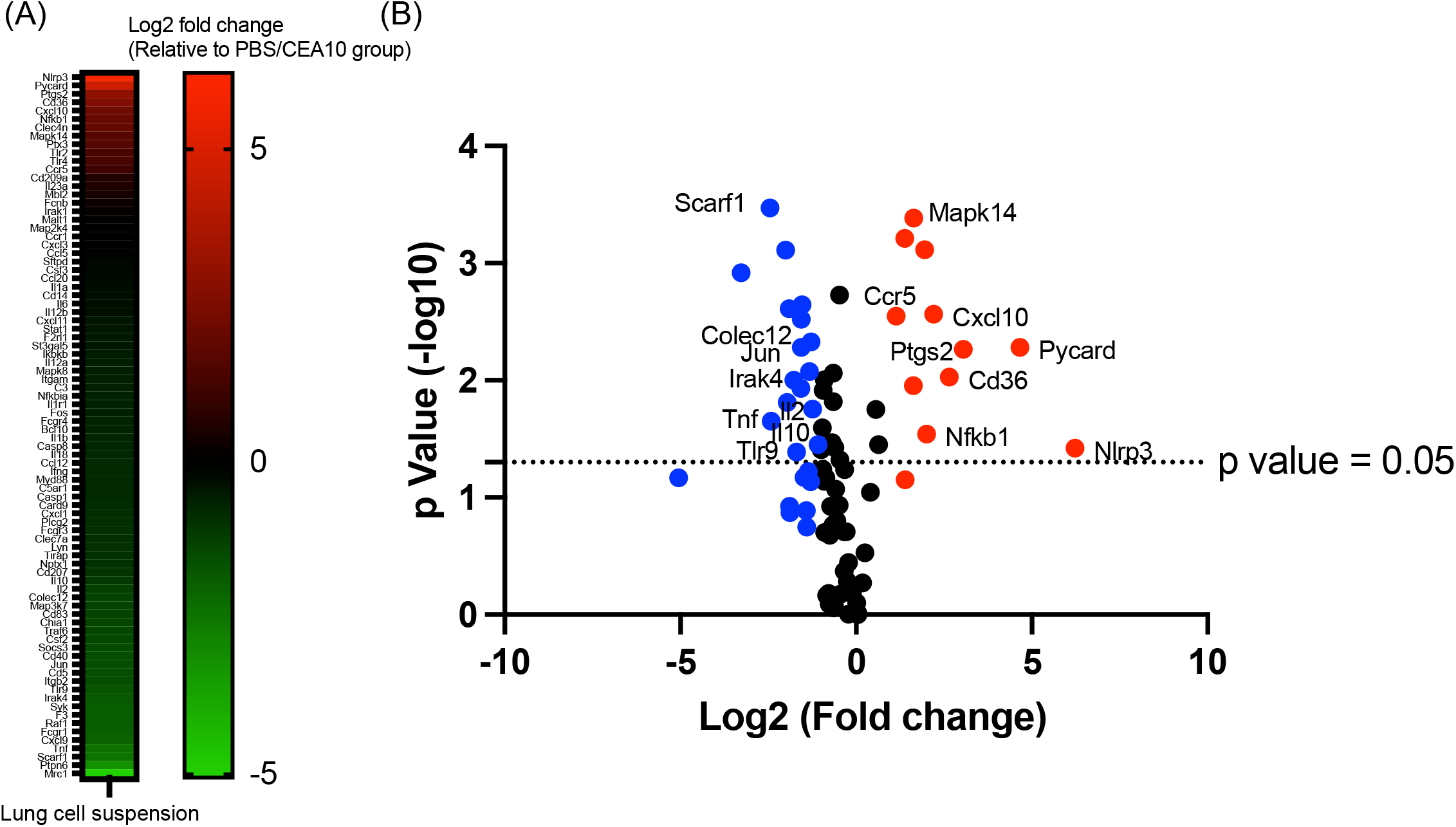
Influenza A virus infection increases transcripts of genes associated with an inflammatory response but reduces transcript levels of known fungal pattern recognition receptors. C57BL/6 mice were inoculated with 100 EID 50 IAV or PBS at Day 0 followed by 3.4*10^7^ CEA10 conidia or PBS at Day 6. Mice were euthanized at 8 hours post CEA10 or PBS inoculation for antifungal gene transcript analysis. Antifungal gene transcript levels were measured by quantitative RT-PCR with RNA from Lung cell suspensions. (A) Increased (red) or Decreased (green) transcript levels of genes associated with antifungal responses in IAV/CEA10 group compared to PBS/CEA10 group represented by the heat map, PBS/CEA10 group: n=3; IAV/CEA10 group: n=3. (B) Volcano plot showed the distribution of fold changes of antifugal gene transcript levels in IAV/CEA10 group compared to PBS/CEA10 group. Genes with increase fold changes >2 were shown in red and genes with decrease fold changes >2 were shown in blue. The p value threshold = 0.05 (Student’s T-test) is indicated by the line in the plot.

## DISCUSSION

Seasonal Influenza infection is a common annual respiratory disease among humans and 1-2% of patients with symptomatic illness require hospitalization in the United States (33). Of those hospitalized, approximately 5-10% progress to ICU admission (34). Patients in the ICU due to severe IAV infections are well known to develop secondary bacterial infections, but in recent years the incidence of secondary fungal infections has been reported with a prevalence of IAPA between 5 to 19% and a mortality rate of ~50% (8, 35–37). Importantly, both immunocompromised and immune competent individuals in the ICU were at risk for developing IAPA (35). Moreover, this does not seem to be isolated to severe respiratory infection with IAV, because recent clinical case reports from the current SARS-CoV2 pandemic suggest that secondary fungal infections are observed in patients with severe COVID19 (38–41). COVID19-associated pulmonary aspergillosis (CAPA) is reported in 20-35% of cases in recent reports from Europe (38). Thus, severe respiratory viral infection is an emerging risk factor for invasive aspergillosis and warrants further mechanistic studies.

Influenza infection can cause epithelial cell damage and leakage, induce antiviral cytokine production, impair further inflammatory cellular recruitment, and impair the phagocytic and antimicrobial activity of macrophages and neutrophils, which all can contribute to the increased risk of developing secondary bacterial infection (12, 13). However, there is a significant knowledge gap in understanding how IAV infection makes the lung environment conducive for fungal growth leading to the development of IAPA. Recently, in a human retrospective observational study Feys *et al* identified a couple breaches in antifungal immunity in patients with IAPA and CAPA which included decreased integrity of the epithelial barrier, decreased antifungal cytokine expression (e.g. IFNγ), and decreased gene expression for pathways involved in fungal phagocytosis and killing mainly mediated by neutrophils (39). One major hurdle to understanding viral-induced pulmonary aspergillosis has been the lack of experimental models to dissect secondary fungal infection. Our work here, together with the model from Robinson and colleagues (17), has established a robust IAV-*Af* superinfection mouse model that can be used for future mechanistic studies. To this end we used our new model to determine whether prior IAV infection created a favorable environment for fungal escape from host antifungal innate immune response, and if so, to experimentally define the host cellular function(s) modulated by the post-IAV lung environment.

Host resistance against *A. fumigatus* can be lost by either quantitative or qualitative defects in the antifungal leukocyte response (40). To begin to investigate how prior IAV infection enhanced susceptibility to *A. fumigatus*, we examined both inflammatory immune cell accumulation in the lungs and their antifungal functions in that environment. Flow cytometry analysis of lung suspensions showed that neutrophil, interstitial macrophage, monocyte, and cDC2 accumulation in the lung parenchyma was similar in IAV-*Af* superinfected and *A. fumigatus* only infected mice (Fig. 2). This is in line with the inferred cellular make-up of the human IAPA bronchoalveolar lavage fluid in human patients with IAPA versus IAV infection along (41). In contrast, Tobin *et al*. previously found that the proportion of neutrophils and alveolar macrophages was decreased in IAV-*Af* superinfected mice compared to *A. fumigatus* only infection (17). The authors did not quantify absolute numbers of inflammatory cells but did note overall increased inflammation in their histological analysis, which could explain our discrepant results. Additionally, our studies utilized the CEA10 strain of *A. fumigatus*, while Tobin *et al* used the ATCC42202 strain (42), which could also drive the differences in our findings since *A. fumigatus* strain heterogeneity alters virulence and host immune responses (43–48). Future studies will examine the role of *A. fumigatus* strain heterogeneity in driving IAPA.

Since we observed no obvious quantitative differences in the innate immune cell response in the IAV-*Af* superinfected mice versus *A. fumigatus* only infection it was likely that the innate immune cells from the IAV-*Af* superinfected mice had functional defects in their antifungal effector response in the post-IAV lung environment. To examine the overall antifungal effector functions of host leukocytes in our murine IAV-*Af* superinfection model, we used the robust *in vivo* FLARE method to determine both conidial uptake and killing by professional phagocytes in the presence or absence of IAV infection (73). We found that neutrophils and monocytes had defects in both conidial uptake and killing in the post-IAV lung environment (Fig. 3). This is in line with what others have observed with regard to leukocyte function in IAV-induced secondary bacterial infections (49, 50). Moreover, this directly supports the transcriptome correlates identified in human patients with IAPA (41).

Innate immune resistance against *A. fumigatus* requires both ROS-dependent and ROS-independent mechanisms for fungal conidial clearance (51, 52). Oxidative stress from host ROS production is known for preventing fungal conidial germination (53). IAV infection can limit ROS production in neutrophils and monocytes after secondary bacterial challenge (54). In contrast, to what was seen with bacterial challenge, we observed no defect in ROS production by both neutrophils and monocytes from IAV-*Af* superinfected mice compared to mice infected with only *A. fumigatus* (Fig. 4). We also noticed that there were more ROS producing neutrophils in IAV single infection, which was consistent with previous IAV infection studies (54, 55). Thus, our data suggest that monocytes and neutrophils from IAV-*Af* superinfected mice maintain robust ROS production in response to secondary fungal challenge, which may be reflective of bacterial infections being highly dependent on TLRs for the induction of ROS, whereas fungal infections are highly dependent on CLRs for the induction of ROS.

ROS-independent conidial clearance in both macrophages and neutrophils requires phagolysosome maturation to create an acidic environment which is necessary for conidial killing (56, 57). A recent study also showed that β-coronavirus infection can lead to lysosome destruction, affect its acidification, and reduce bacterial clearance in macrophage (58). To assess phagolysosome maturation in the neutrophils and monocytes within the IAV-*Af* superinfected lungs, we used pHrodo-Zymosan staining. Both neutrophils and monocytes displayed a significant reduction in pHrodo-Zymosan signal following both IAV infection alone or IAV-*Af* superinfection (Fig. 5), which is supportive of impaired or slowed phagolysosome maturation in the post-IAV lung environment. In addition to the phagolysosome maturation level, we also examined the percentage of cells that are still able to uptake and send the pHrodo-Zymosan into mature phagolysosomes through the percentage of pHrodo-Zymosan positive cells in neutrophil and monocyte populations. Interestingly, we observed a significant reduction of monocytes that can response and send pHrodo-Zymosan into mature phagolysosome in the post-IAV environment. This suggests that the defects are in both PAMP responsiveness and phagolysosome maturation in the post-IAV environment for monocytes (Fig. S8B). However, we did not detect a significant reduction in percentage of pHrodo-zymosan^(+)^ in neutrophils from IAV infected mice, indicating potential differences between neutrophils and monocytes in the post-IAV environment. Instead, the significant reduction of pHrodo-Zymosan uptake in both the *A. fumigatus* single infection and IAV-*Af* superinfection groups might be due to potential neutrophil exhaustion during fungal infection since the pHrodo-Zymosan uptake requires the recognition and binding to the dectin-1 receptor (Fig. S8A). Still, we can detect a decrease in mature phagolysosome and conidial killing in the pHrodo-Zymosan^(+)^ neutrophils, suggesting that the phenotype is not contributed by cell exhaustion post-IAV infection (Fig. 6C and 6E). In summary, our data revealed that the post-IAV environment does not inhibit immune cell recruitment or ROS production in the face of a highly virulent *A. fumigatus* strain, but specifically reduces phagolysosome maturation in neutrophils and monocytes, which can lead to conidial escape from the host innate immunity during fungal infection.

A major question remaining from our study is how the post-IAV lung environment impairs phagolysosome maturation. Susceptibility to secondary fungal infections are warranted. To investigate the upstream signal that drives defective fungal clearance in the IAV-*Af* superinfection mice in our model, we examined antifungal gene expression in isolated immune cells. Based on viral-bacterial superinfection literature, we expected to observe an antiinflammatory expression profile after IAV infection and during early fungal infection (58–61). However, we observed that immune cells from IAV-*Af* superinfection mice displayed increased inflammatory gene transcript levels and decreased anti-inflammatory cytokine gene transcript levels (Fig. 7), indicating enhanced immune responses during early fungal infection. This result may be due to the increase in fungal burden in the superinfection group. Interestingly, we found reduced expression of three PRRs (*Tlr9, Scarf1, Colec12*) (Fig. 7). TLR9 is known to be recruited to the *A. fumigatus-containing* phagosome and contribution to fungal induced immune cell activation (59, 60). On the other hand, TLR9 trafficking to the phagosome is mediated by Dectin-1 signaling and phagolysosome maturation and acidification(61, 62). Although we did not observe a difference in *Clec7a* transcript levels in the IAV-*Af* superinfection mice compared to *A. fumigatus* only infection group, the decreased *Tlr9* expression and reduced phagosome acidification in our IAV-*Af* superinfection model suggests impaired TLR9 activation and signaling. In addition to decreased *Tlr9* expression, we also observed decreased *Irak4* expression (Fig. 7), which is a component of the myddosome needed for TLR-dependent signaling (63). Therefore, it is likely that reduction of TLR9 signaling contributes to defects in phagolysosome maturation in neutrophils and monocytes, or vice versa.

LC3-associated phagocytosis (LAP) is known to be associated with more rapid phagolysosome maturation (64). LAP is known to be essential for host resistance against *A. fumigatus* (65, 66). In human IAPA and CAPA patients, genes (*MAP1LC3B* and *SQSTM1*) related to LC3-associated phagocytosis (LAP) were decreased while *CDC20*, the gene involved in LC3 degradation was upregulated in the superinfection group comparing to viral single infection (41). Following CpG oligonucleotide activation of TLR9, LC3 can be recruited to the signaling endosome and mediate IKKα recruitment that leads to TRAF3 and IRF7 activation Additionally, IFNγ enhances LAP following *A. fumigatus* challenge, but in humans with IAPA the levels of IFNγ are diminished to those with IAV only (41). Thus, we hypothesize that reducing TLR9 signaling in our IAV-*Af* superinfection model contributes to a decrease in LAP, further linking phagolysosome maturation and *A. fumigatus* clearance (67). This link between phagosome maturation and effective fungal clearance is well supported by mechanistic studies in other fungal pathogens, where phagolysosome maturation is key to clearance of infection (68–70). Still, further investigation of the role of LAP components specifically and contributions of TLR9 signaling in our IAV-*Af* superinfection model is required to address this possibility and to assess whether therapeutically targeting this pathway could restore host resistance against *A. fumigatus* after IAV infection.

In conclusion, we found that the post-IAV lung environment negatively affects phagolysosome maturation, which corresponds reduced antifungal killing by both neutrophils and monocytes. This reduction in fungal killing leads to increased fungal germination, fungal growth, and eventual establishment of IAPA. Further studies are needed to investigate the upstream signals altering phagolysosome maturation in the presence of IAV. More information could be leveraged therapeutically to restore antifungal activity of impaired neutrophils and monocytes after IAV infection, to enhancing fungal clearance and clinical outcomes in IAPA patients. The findings from this study are also more broadly applicable, as recent clinical case reports have revealed that patients with severe COVID-19 can develop a secondary infection with *A. fumigatus* and these patients had worse clinical outcomes and higher mortality. It remains to be determined what the underlying mechanisms are for SARS-CoV2 induced secondary *A. fumigatus* infection.

## MATERIALS AND METHODS

### Animal Inoculation

C57BL/6J mice between 8-10 weeks old were purchased from Jackson Laboratories. Mice were housed in autoclaved cages at ≤ 4 mice per cage with a supply of HEPA-filtered air and water. Only mice with a weight under 22 grams were selected for all experiments. The stock of Influenza A/PR/8/34 H1N1 was purchased from Charles River and the titer of the virus was quantified by egg infectious dose (EID50). *A. fumigatus* strain CEA10 (also called CBS144.89) was grown on a 1% glucose minimal media (GMM) plate for 3 days at 37 °C. The conidia were collected in 0.01% Tween 20 and washed 3 times with sterile PBS. Mice were infected with 100 EID50 of Influenza A/PR/8/34 H1N1 (in 50 μl sterile PBS) or control PBS by intranasal instillation under isoflurane anesthesia. After 6 days of viral infection, mice were infected with 3.4×10^7^ CEA10 (in 100 μl sterile PBS) or control PBS by oropharyngeal instillation. Mice were then euthanized between 8 to 48 hours *post-Aspergillus* challenge. Animals were monitored daily for disease symptoms and we carried out our animal studies in strict accordance with the recommendations in the Guide for the Care and Use of Laboratory Animals. The animal experimental protocol 00002167 was approved by the Institutional Animal Care and Use Committee (IACUC) at Dartmouth College.

### RNA Preparation and Pathogen Quantification

Mice were inoculated with either PBS, Influenza A/PR/8/34 H1N1, or CEA10 as described previously and the lungs were removed at euthanasia for RNA extraction. Lungs were flash frozen, lyophilized and homogenized with glass beads using Mini Bead Beater (BioSpec Products Inc, Bartlesville, OK) and resuspended in Trizol reagent (Thomas Scientific) and chloroform to extract RNA according to manufacturer’s instruction. 5 μg of RNA was treated with Ambion Turbo DNase (Life Technologies) according to the manufacturer’s instruction. I μg of DNase treated RNA was further processing with QuantiTech Reverse Transcription kit with additional 0.5 ng of Random Decamer. The RNA amounts were normalized to *Gapdh* for fungal and viral burden. Primers used were: Forward primer for murine GAPDH: 5’-TCATCCCAGAGCTGAACG-3’, reverse primer for murine GAPDH: 5’-GGGAGTTGCTGTTGAAGTC-3’. The fungal burden was measured by quantitative RT-PCR on *Aspergillus fumigatus* 18s rRNA. Primers used were: Forward primer for fungal burden: 5’- GGCCCTTAAATAGCCCGGT -3’, reverse primer for fungal burden: 5’- TGAGCCGATAGTCCCCCTAA-3’, Taqman probe: AGCCAGCGGCCCGCAAATG. The viral load was measured by quantitative RT-PCR on viral matrix protein (17). Primers used were: Forward primer for viral load: 5’- GGACTGCAGCGTAGACGCTT -3’, reverse primer for viral load: 5’- CATCCTGTTGTATATGAGGCCCAT -3’, PrimeTime probe: 5’-/56-FAM/CTCAGTTAT/ZEN/TCTGCTGGTGCACTTGCCA/ 3IABkFQ/-3’.

### Histology Staining

Mice were inoculated with either PBS, Influenza A/PR/8/34 H1N1 or CEA10 as described previously and euthanized at 48 hours post fungal inoculation. After euthanasia, cannulas were inserted into the trachea and the lungs were removed from the body cavity. The lungs were inflated and immersed in 10% buffered formalin phosphate for 24 hours and stored in 70% ethanol until embedding. Paraffin-embedded sections were stained with hematoxylin and eosin (H&E) to observe inflammation and Grocott-Gomori methenamine silver (GMS) to observe fungi. The images of H&E and GMS slides were analyzed microscopically with a Zeiss Axioplan 2 imaging microscope (Carl Zeiss Microimaging, Inc., Thornwood, NY) fitted with a QImaging Retiga-SRV Fast 1394 RGB camera.

### Flow Cytometry: Lung Cellularity and Fluorescence Aspergillus REporter (FLARE) Analysis

For the FLARE experiments, the mRFP expressed CEA10 conidia were generated by ectopically insertion of *gpdA*-driven mRFP construct with *ptrA* gene as selection marker. To generate FLARE conidia, mRFP expressed conidia were collected and labeled with Alexa633 as described previously (71). Mice were inoculated with either PBS, Influenza A/PR/8/34 H1N1 or FLARE conidia as described previously and euthanized at 36 hours post fungal inoculation. To harvest single cell suspension from mice lungs, the whole lungs were minced and digested in buffer containing 2.2 mg/ml Collagenase type IV (Worthington), 1 U/ml DNase1 (New England Biotech) and 5% FBS at 37 °C for 45 min. The digested samples were passed through 18-gauge needle, incubated in RBC Lysis buffer (eBioScience), neutralized in PBS, passed through 100 um filter and counted. The antibodies used for the Flow cytometry analysis on different populations as the following: For neutrophil population, lung cells were stained with Survival dye (efluor780, eBioScience), CD45 (Pacific Orange, Invitrogen), CD64 (BV421, BioLegend), Ly6G (FITC, BioLegend), CD11b (PercpCy5.5, BioLegend). For macrophage/monocyte population, lung cells were stained with Survival dye (efluor780, eBioScience), IA/IE (MHCII) (BV605, BioLegend), SiglecF (BV421, BD BioScience), Ly6G (FITC, BioLegend), CD103 (FITC, BioLegend), CD11b (PercpCy5.5, BioLegend), CD64 (PECy7, BioLegend). For DC populations, lung cells were stained with Survival dye (efluor780, eBioScience), IA/IE (MHCII) (BV605, BioLegend), CD11b (Pacific Blue, BioLegend), CD103 (FITC, BioLegend), CD317 (PercpCy5.5, BioLegend), CD11c (PECy7, BioLegend). The gating strategy of each cell populations are indicated in Fig. S1-S3. The data were collected by Beckman Coulter Cytoflex S and analyzed with FlowJo version 10.8.1. To quantify fungal colony forming units (CFUs) of intracellular conidia, lung cells in the single cell suspension were further homogenized with glass beads using Mini Bead Beater (BioSpec Products Inc, Bartlesville, OK) and resuspended in PBS. The samples were diluted 1:100 and then plated on the ½ Sabouraud dextrose agar plates, incubated overnight, and counted for CFU.

### Immune Cell Function: Intracellular ROS Production and Phagolysosome Maturation

Mice were inoculated with either PBS, Influenza A/PR/8/34 H1N1 or CEA10 conidia as described previously and euthanized at 8 hours post fungal inoculation. The single cell suspensions were harvested as described previously. For measurement of intracellular ROS, lung cells from the single cell suspensions were incubated with 1 μM of CM-H2DCFDA (5-(and-6)-chloromethyl-2’,7’-dichlorodihydrofluorescein diacetate, Thermo) at 37 °C for 30 minutes according to manufacturer’s instruction. For the measurement of phagolysosome maturation, lung cells were incubated with 0.05 mg/ml of pHrodo Green Zymosan Bioparticles (‘pHrodo-Zymosan’, Invitrogen) at 37 °C for 2 hours according to manufacturer’s instruction. CM-H2DCFDA or pHrodo-Zymosan stained lung cells were then stained with Survival dye (efluor780, eBioScience), IA/IE (MHCII) (BV605, BioLegend), SiglecF (BV421, BD BioScience), Ly6G (PE, BioLegend), CD11b (PercpCy5.5, BioLegend), CD64 (PECy7, BioLegend) for the gating of neutrophils and monocytes. The gating strategy of each cell populations were indicated in Fig. S6. For confirming intracellular localization of signals by microscopy and verifying flow cytometry results in Fig. 6, mice were inoculated with CEA10-mRFP FLARE conidia and lung cells were incubated with 0.05 mg/ml of pHrodo-Zymosan at 37 °C for 2 hours and stained with Survival dye (efluor780, eBioScience) and Ly6G (Pb, BioLegend) or Ly6G (Pb, BioLegend) alone. Images were acquired using an Andor W1 spinning disk confocal microscope, mounted with a Nikon Eclipse Ti inverted microscope stand. Lasers with wavelengths of 405 nm (Ly6G, Pb, BioLegend), 488 nm (pHrodo-Zymosan, Invitrogen), 561 nm (CEA10-mRFP) and 633 nm (Alexafluor 633 labeled conidia, Invitrogen) were used for excitation. Images were viewed using Fiji Software, and were used to visualize flow cytometry results. Flow Cytometry data were collected by Beckman Coulter Cytoflex S and analyzed with FlowJo version 10.8.1.

### RNA preparation and antifungal gene expression evaluation

Mice were inoculated with either PBS, Influenza A/PR/8/34 H1N1 or CEA10 conidia as described previously and euthanized at 8 hours post fungal inoculation. The single cell suspensions were harvested as described previously. Following the cell isolation, the cells were lysis and RNA was collected by RNeasy kit according to manufacturer’s instruction (Qiagen). 0.5 μg of RNA was further processing with RT^2^ First Strand Kit (Qiagen). cDNA was mixed with RT2 SYBR^®^ Green Mastermix (Qiagen) and loaded to RT^2^ Profiler PCR Arrays containing primers for antifungal genes (PAMM-147ZD; table S2; Qiagen). The RNA load was normalized with *Actb, Gapdh* and *Hsp90ab1*.

### Statistical Analysis

All statistical analyses were performed with Prism 9.3.0 software (GraphPad Software Inc., San Diego, CA). The Log-rank test and Gehan-Breslow-Wilcoxon test were performed for statistical analysis of the survival curve. For animal experiments, nonparametric analyses were performed (Kruskal-Wallis, Dunn’s multiple comparisons; Mann-Whitney, single comparisons). All error bars represent standard deviations. NS P>0.05; * P≤0.05; ** P≤0.01; *** P≤0.001; **** P≤0.0001.

## ACKNOWLEDGMENTS

Special thanks to Center for Comparative Medicine and Research (CCMR) staff Eric Dufour for helping with the animal experiments. Flow Cytometry data were collected with technical assistance and resources from the Immune Monitoring and Flow Cytometry Resource (IMFCSR) at the Norris Cotton Cancer Center at Dartmouth. This project was supported by NIH/NIAID grant R21-AI152019 (JJO), NIH/NIAID R01AI39632 (RAC), and a Cystic Fibrosis Foundation Research Development Grant (STANTO15RO). MSG was supported by the Dartmouth College Immunology Training Program (NIH/NIAID T32 AI007363). We also thank the Imaging Facility at Dartmouth (Ann Lavanway) and the Biomolecular Targeting Core (P20-GM113132) for use of equipment.

## Supplemental Material Legends

Fig. S1 Gating strategy for Neutrophils. The neutrophils were identified as CD45^(+)^Ly6G^(+)^CD11b^(+)^cells and free conidia as FSC^(low)^SSC^(low)^ cells.

Fig. S2 Gating strategy for Macrophages/Monocytes. The alveolar macrophages were identified as Ly6G^(-)^CD103^(-)^SiglecF^(+)^CD11b^(+)^ cells, interstitial macrophages as Ly6G^(-)^CD103^(-)^SiglecF^(-)^CD11b^(hi)^CD64^(+)^ cells and monocytes as Ly6G^(-)^CD103^(-)^SiglecF^(-)^CD11b^(hi)^CD64^(-)^MHCII^(-)^ cells.

Fig. S3 Gating strategy for DCs. The CD103^(+)^ cDC1 were identified as MHCII^(+)^CD11c^(+)^CD11b^(-)^CD103^(+)^cells, CD11b^(+)^ cDC2 were identified as MHCII^(hi)^CD11c^(hi)^CD103^(-)^CD11b^(+)^ cells and pDC were identified as MHCII^(+)^CD11c^(+)^CD11b^(-)^CD103^(-)^CD317^(+)^ cells.

Fig. S4 Influenza A virus infection does not affect immune cell composition during IA. C57BL/6 mice were inoculated with 100 EID 50 IAV or PBS at Day 0 followed by 3.4*10^7^ CEA10 conidia or PBS at Day 6. Mice were euthanized at 36 hours post CEA10 or PBS inoculation for lung cellularity experiments. Three independent experiments were performed and data are shown as the combined results (PBS/PBS group: n=7; IAV/PBS group: n=8; PBS/CEA10 group: n=22; IAV/CEA10 group: n=22). The percentage of lung cells was acquired by flow cytometry as indicated as: (A) neutrophils (CD45^(+)^Ly6G^(+)^CD11b^(+)^), (B) alveolar macrophages (Ly6G^(-)^CD103^(-)^SiglecF^(+)^CD11b^(+)^), (C) interstitial macrophages (Ly6G^(-)^CD103^(-)^SiglecF^(-)^CD11b^(hi)^CD64^(+)^), (D) monocytes (Ly6G^(-)^CD103^(-)^SiglecF^(-)^CD11b^(hi)^CD64^(-)^MHCII^(-)^), (E) CD103^(+)^ cDC1 (MHCII^(+)^CD11c^(+)^CD11b^(-)^CD103^(+)^), (F) CD11b^(+)^ cDC2 (MHCII^(hi)^CD11c^(hi)^CD103^(-)^CD11b^(+)^), (G) pDC (MHCII^(+)^CD11c^(+)^CD11b^(-)^CD103^(-)^CD317^(+)^). Kruskal-Wallis with Dunn’s multiple comparisons was performed for statistical analyses. All error bars represent standard deviations. NS P>0.05; * P≤0.05; ** P≤0.01; *** P≤0.001; **** P≤0.0001.

Fig. S5 FLARE results of lung phagocytes. The gating for FLARE experiments in the (A) neutrophils and free conidia, (B) alveolar macrophages, interstitial macrophages and monocytes, (C) CD11b^(+)^ cDC2. R1 denotes phagocytes containing live conidia. R2 denotes phagocytes containing killed conidia. (R1+R2) indicates conidial uptake of phagocytes and (R1/(R1+R2)) indicates conidial viability in the phagocytes.

Fig. S6 Gating strategy for pHrodo/ROS experiments. The neutrophils were identified as Ly6G^(+)^cells and monocytes as Ly6G^(-)^CD103^(-)^SiglecF^(-)^CD11b^(hi)^CD64^(-)^MHCII^(-)^ cells. Representative dot plots were from pHrodo-Zymosan staining experiment.

Fig. S7 Post influenza environment does not reduce ROS productive Neutrophils and Monocytes. C57BL/6 mice were inoculated with 100 EID 50 IAV or PBS at Day 0 followed by 3.4*10^7^ CEA10 conidia or PBS at Day 6. Mice were euthanized at 8 hours post CEA10 or PBS inoculation. Lung cell suspensions were stained with CM-H2DCFDA for 30 minutes and then stained for neutrophils and monocytes. The percentage of ROS producing cells was shown as the percentage of cells with positive signal from CM-H2DCFDA staining in (A) neutrophils (Ly6G^(+)^), (B) monocytes (Ly6G^(-)^CD103^(-)^SiglecF^(-)^CD11b^(hi)^CD64^(-)^MHCII^(-)^). Two independent experiments were performed and data are shown as the combined results (PBS/PBS group: n=8; IAV/PBS group: n=8; PBS/CEA10 group: n=12; IAV/CEA10 group: n=11). Kruskal-Wallis with Dunn’s multiple comparisons was performed. All error bars represent standard deviations. NS P>0.05; * P≤0.05; ** P≤0.01; *** P≤0.001; **** P≤0.0001.

Fig. S8 Post influenza environment hinders the phagocytosis of Monocytes. C57BL/6 mice were inoculated with 100 EID 50 IAV or PBS at Day 0 followed by 3.4*10^7^ CEA10 conidia or PBS at Day 6. Mice were euthanized at 8 hours post CEA10 or PBS inoculation for phagolysosome maturation analysis. Lung cell suspensions were incubated with pHrodo-Zymosan for 2 hours and then stained for neutrophils and monocytes. The percentage of active cells with mature phagolysosome was measure by the percentage of cells with positive signal from the color change of pHrodo-Zymosan in (A) neutrophils (Ly6G^(+)^), (B) monocytes (Ly6G^(-)^SiglecF^(-)^CD11b^(hi)^CD64^(-)^MHCII^(-)^). Two independent experiments were performed and data are shown as the combined results (PBS/PBS group: n=8; IAV/PBS group: n=8; PBS/CEA10 group: n=12; IAV/CEA10 group: n=11). Kruskal-Wallis with Dunn’s multiple comparisons was performed for statistical analyses. All error bars represent standard deviations. NS P>0.05; * P≤0.05; ** P≤0.01; *** P≤0.001; **** P≤0.0001.

Table S1 Staining panels for Flow Cytometry. Antibody lists for (A) neutrophils and free conidia. (B) alveolar macrophages, interstitial macrophages and monocytes. (C) CD103^(+)^ cDC1, CD11b^(+)^cDC2 and pDC. (D) neutrophils and monocytes in the pHrodo and ROS staining experiments. Cellular markers for Flow Cytometry. Cellular markers for (E) neutrophils. (F) alveolar macrophages, interstitial macrophages and monocytes. (G) CD103^(+)^ cDC1, CD11b^(+)^ cDC2 and pDC. (H) neutrophils and monocytes in the pHrodo and ROS staining experiments.

Table S2 Gene list of antifungal RT2 Profiler PCR Arrays. cDNA of mice from PBS/CEA10 (n=3) and IAV/CEA10 groups (n=3) was added to the plates containing primer sets in the table. The RNA load was normalized with *Actb, Gapdh* and *Hsp90ab1*. The gene expression was compared by IAV/CEA10 group over PBS/CEA10 group. Genes with increase fold changes >2 were shown in red and genes with decrease fold changes >2 were shown in blue.

